# Sustained dechlorination of vinyl chloride to ethene in *Dehalococcoides*-enriched cultures grown without addition of exogenous vitamins and at low pH

**DOI:** 10.1101/612242

**Authors:** Luz A. Puentes Jácome, Po-Hsiang Wang, Olivia Molenda, Yi Xuan (Jine-Jine) Li, M. Ahsanul Islam, Elizabeth A. Edwards

## Abstract

Trichloroethene (TCE) is a ubiquitous groundwater pollutant. Successful TCE bioremediation has been demonstrated at field sites using specialized microbial consortia harboring TCE-respiring *Dehaloccocoides* whose growth is cobalamin (vitamin B_12_)-dependent. Bioaugmentation cultures grown ex situ with ample exogenous vitamins in the medium and at neutral pH may become vitamin-limited or inhibited by acidic pH once injected into field sites, resulting in incomplete TCE dechlorination and accumulation of more toxic vinyl chloride (VC). Here, we report growth of the *Dehalococcoides*-containing bioaugmentation culture KB-1 in a TCE-amended mineral medium devoid of vitamins and in a VC-amended mineral medium at low pH (6.0 and 5.5). In cultures grown without exogenous vitamins or cobalamin, *Acetobacterium*, which can synthesize 5,6-dimethylbenzimidazole (DMB), the lower ligand of cobalamin, and *Sporomusa* are the dominant acetogens. At neutral pH, a growing *Acetobacterium* population supports complete TCE dechlorination by *Dehalococcoides* at millimolar levels with a substantial increase in the amount of measured cobalamin (~20-fold). Sustained dechlorination of VC to ethene was achieved at a pH as low as 5.5, yet at low pH *Acetobacterium* is less abundant, potentially affecting the production of DMB and/or cobalamin. However, dechlorination activity at very low pH (< 5.0) was not stimulated by DMB supplementation, but was restored by raising pH to neutral. Assays in cell extracts revealed that vinyl chloride reductase (VcrA) activity declines significantly below pH 6.0 and is undetectable below pH 5.0. This study highlights the roles of and interplay between vitamin-producing populations and pH in microbial dechlorinating communities, and their importance for successful chlorinated ethenes bioremediation at field sites.

## INTRODUCTION

*Dehalococcoides* and *Dehalogenimonas* spp., the only known bacteria capable of reductive dechlorination of VC to non-toxic ethene,^1–4^ require H2 as electron donor and conserve energy through organohalide respiration. Organohalide respiration depends on reductive dehalogenases (RDases) that harbour a cobamide as a prosthetic group. However, *Dehalococcoides* spp. cannot synthesize their own cobamide and thus import the vitamin from their surroundings. *Dehalococcoides*-containing cultures and isolates are typically grown in the presence of cobamides like cobalamin (vitamin B_12_).^5, 6^ Cobamides are composed of a tetrapyrrole ring, a central chelated cobalt ion, two axial ligands (upper and lower ligands), an aliphatic side chain and a phosphodiester bond which links the tetrapyrrole ring and the α-ribazole (i.e., the ribonucleoside with the lower axial ligand).^7, 8^ The best known cobamide is cobalamin which contains5,6-dimethylbenzimidazole (DMB) as the lower axial ligand. A recent study^9^, using *Dehalococcoides mccartyi* strain GT, demonstrated that the vinyl chloride reductase (VcrA)-mediated dechlorination of VC to ethene is achieved only when the lower ligand of the cobamide is DMB or 5-methylbenzimidazole.^9^ The cobamide requirement of *Dehalococcoides* may be satisfied via direct cobalamin addition or via DMB salvage from microbial community partners and subsequent cobamide remodeling.^10, 11^ Currently, only prokaryotes are known to produce cobamides with 17 different lower axial ligand structures,^7^ and generally, only one type is produced by a single family. For example, members of the family *Veillonellaceae*, such as *Sporomusa*, produce cobamides with a phenolic lower base^10, 12^ while members of the family *Eubacteriaceae*, such as *Acetobacterium woodii* and *Eubacterium limosum*, exclusively produce cobamides with a DMB lower ligand.^13, 14^

The commercialized bioaugmentation enrichment culture KB-1^®^ dechlorinates chlorinated ethenes, including the dry-cleaning solvent tetrachloroethene (PCE) and the industrial solvent trichloroethene (TCE). In the laboratory, the original KB-1 parent culture (TCE/M_1998_parent, Figure S1) is maintained with TCE (electron acceptor) and methanol (electron donor). This culture is dominated by several distinct strains of *Dehalococcoides* ^15, 16^ and also contains methanogens and acetogens. The historically dominant acetogens in the parent culture belong to the genera *Sporomusa* and *Acetobacterium*.^3, 17, 18^ The most abundant RDase is VcrA^19^ whose^19^ activity is dependent on cobalamin availability. Hence, dechlorinating enrichment cultures such as KB-1 are typically grown in a medium amended with cobalamin and other vitamins to support steady dechlorination.

Another important factor influencing the performance of dechlorinating cultures is pH. Extensive dechlorination generates HCl which can lead to a drop in pH as the buffering capacity of the medium is exhausted. A drop in pH to below pH 6.0 causes a significant reduction in dechlorination rates and extent.^20–23^ Lower pH may impact the dechlorinators directly, or may impact associated fermenting and cobalamin-producing organisms in the culture. *Dehalococcoides* populations recently studied by Yang *et al*.^24^ could not sustain reductive dechlorination at pH 5.5, yet the reason for the loss of dechlorination activity remains unclear. In addition, the enrichment cultures grown at pH 5.5, in which partial dechlorination of TCE to *cis*-dichloroethene (*cis*-DCE) was observed, were shown to harbour dechlorinating *Sulfurospirrillum*, but not *Dehalococcoides* populations.^24^

Addition of vitamins and cobamides to a field site can be very expensive; vitamin B_12_ (cobalamin) for field site remediation applications is sold at around 1,800 USD per 100 g.^25^ Similarly, pH adjustment at field sites can also be difficult and costly. The first objective of this study was to determine if a TCE-dechlorinating enrichment culture like KB-1, that stoichiometrically dechlorinates TCE to ethene, could be successfully maintained without exogenous vitamins. We have established a subculture of KB-1 that has sustained dechlorination of TCE to ethene without addition of vitamins for over six years, indicating an adequate endogenous supply of vitamins in the mixed culture. The second objective of this study was to determine if *Dehalococcoides* populations can sustainably dechlorinate VC to ethene at progressively lower pH, and to identify the reasons for decreased activity at low pH. Following progressive adaptation, sustained VC dechlorination to ethene was observed in KB-1 cultures maintained at pH ~ 6.0 and even at pH ~ 5.5, but not observed at lower pH. We determined that the rate of VC dechlorination to ethene by *Dehalococcoides* in KB-1 subcultures at pH below 5.0 was not limited by DMB or cobalamin availability, but rather by a direct effect of pH on the VcrA-mediated dechlorination activity. The impact of these results is discussed in the context of bioremediation practice.

## MATERIALS AND METHODS

### Microorganisms and chemicals

Chemical reagents were purchased through Sigma-Aldrich Canada (Oakville, ON, Canada), Fischer Scientific Canada (Ottawa, ON, Canada), and BioShop (Burlignton, ON, Canada) at the highest purity available. Gases were purchased from Praxair (Mississauga, ON, Canada). P-cresol-cobamide^12^ and the pure culture of *Sporomusa* sp. KB-1 were provided by Prof. Frank E. Löffler (University of Tennessee Knoxville, USA).

### Analytical Procedures

Sampling, DNA extraction, quantitative PCR (qPCR), 16S rRNA gene amplicon sequencing, and cobamide extraction and measurement protocols are detailed in the Supplementary Information (SI).

### Medium preparation

A bicarbonate-buffered FeS-reduced mineral medium^26, 27^ was used to maintain enrichment cultures and set up new experiments. The medium was either prepared without any vitamins, or with addition of one of two vitamin stock solutions: 1) a complete stock containing 11 vitamins (as described in Edwards and Grbić-Galić ^28^), or 2) the same stock prepared without any vitamin B_12_. The medium was autoclaved and purged while cooling with a gas mix containing 80% N2 and 20% CO_2_. The pH of the medium was 6.9 ± 0.1. If required, the pH was adjusted using a 6N HCl solution. Except for the *Sporomusa* growth inhibition assay, methanol was always used as the sole electron donor and carbon source. TCE was fed in solution with methanol, and VC (pure gas) was fed using a gastight syringe. The TCE/methanol solution was prepared at a ratio of 5:1 electron equivalents (eeq) of donor to acceptor (Methanol 6 eeq/mol; TCE to ethene: 6 eeq/mol).

### Vitamin-free enrichment cultures

Preparation of the initial TCE-dechlorinating enrichment cultures grown and maintained without the addition of exogenous vitamins is described in the doctoral thesis of Islam (2014).^29^ Briefly, three parallel sets of triplicate cultures (100 mL) were started with inoculum from a 4 L TCE-to-ethene-dechlorinating culture derived from the KB-1 parent culture TCE/M_1998_parent (T3MP1), i.e., an enrichment culture maintained since 1998 with TCE as acceptor and methanol (M) as donor. Cells were triple-washed in medium devoid of vitamins, i.e., centrifuged (8000 rpm, 4°C), resuspended, and centrifuged again after discarding the supernatant before inoculation. The first set received medium containing the regular set of vitamins (controls), the second set received medium with all vitamins except cobalamin, and the third set received medium that was completely vitamin-free. The cultures were routinely reamended with donor and acceptor, and periodically transferred into the same three medium formulations. Following the experiments described in Islam (2014)^29^, the cultures (i.e., triplicate 100 mL cultures grown at the three different media formulations described above) were combined and scaled-up to 700 mL and are referred to as TCE/M_Vit(+), TCE/M_Vit(−), and TCE/M_B12(−) (Figure S1).

### Low pH Enrichment cultures

A long term pH adaptation experiment is described in the Master’s thesis of Li (2012)^30^, in which transfers from TCE/M_1998_parent (T3MP1) (Figure S2) were grown over multiple feedings in media at pH 7 (control), pH 6.0, and pH 5.5. In these cultures, the electron acceptor added was VC rather than TCE to maintain specific selection pressure for *Dehalococcoides*. Triplicate enrichment cultures (50 mL) kept at the three pH conditions were maintained with regular VC/methanol feedings for about two and a half years, prior to combining triplicates and scaling up to three 500 mL cultures. These scaled-up cultures are referred to as VC/M_pH7, VC/M_pH6, and VC/M_pH5.5 (Figure S2).

### Growth experiments

Two major growth experiments were performed to test the impacts of pH and B12, one using 10% dilution transfers set up at pH 7.0 and 5.5 (amended with either TCE or VC) and another using 1% dilution transfers (amended with TCE) (see Figure S1 for experimental flow). All experiments were set up in an anaerobic chamber (Coy) with a 10% CO_2_, 10% H_2_, and 80% N_2_ atmosphere. For the 10% dilution transfers at pH 7.0 and 5.5 (Figure S1), inoculum from the TCE/M_B12(−) enrichment culture (10 mL for TCE dechlorination assays and 5 mL for VC dechlorination assays) was added to autoclaved 160-mL narrow-necked glass serum bottles containing defined mineral medium without cobalamin (90 mL for TCE dechlorination assays and 45 mL for VC dechlorination assays). Subsequently, the serum bottles were sealed with rubber stoppers and crimped. The electron donor (methanol) and electron acceptor (TCE or VC) were added in the same way as for the enrichment cultures, at a ratio of 5 to 1 electron equivalents of donor to acceptor, unless otherwise specified. If necessary, additional electron donor was added to ensure complete dechlorination. Once the TCE transfers set up at pH 5.5 had partially dechlorinated the added TCE to ethene, they were subsequently aliquoted to six 10 mL glass vials to a final volume of 5 mL each, sealed with rubber-stoppers and crimped. DMB (10 μM) was added to two bottles and the pH was adjusted to 7.0 in another two bottles using a saturated bicarbonate solution (~ 1.3 M). The remaining two bottles were kept as controls at pH 5.5. The growth of *Sporomusa* and *Acetobacterium* in the absence of TCE was investigated in 10% dilution transfers from the TCE/M_B12(−) culture that were fed methanol only (Figure S1).

The 1% dilution transfer experiments (Figure S1) were initiated to minimize cobalamin and DMB carry-over. In these experiments, the conditions were similar to that of the 10% dilution transfers, except that the inoculum (1 mL) from the TCE/M_Vit(−) was first pelleted down, washed twice with sterilized defined (cobalamin-free) medium, resuspended in 1 mL of same cobalamin-free medium, and then added to serum bottles containing 99 mL of cobalamin-free medium.

### Cell-extract dehalogenase activity assay

The cell-extract dehalogenase activity assay was performed following previously established protocols under anaerobic conditions.^31^ All reagents were pre-reduced or kept anaerobically before the assays. Briefly, 100 mL of the TCE/M_B12(−) culture was harvested and pelleted down by centrifugation at 9,000 x g for 20 min at 4°C under anaerobic conditions. The supernatant was removed, the pellets were resuspended in 0.3 mL of 100 mM Tris-HCl buffer (pH 7.5) containing 12.5 mM NaCl, 2.5% glycerol, and 1% digitonin (detergent). The suspensions were loaded into a 2 mL O-ring-capped plastic microcentrifuge tube containing 50 mg of glass beads (≤ 106 μm). To lyse the cells, the tube was vortexed at maximum intensity for 2 min and incubated on ice for 1 min. The step was repeated 4 times. The cell lysate was then centrifuged at 16,000 x g for 15 min at 4°C to remove cell debris, beads, and unbroken cells, and the supernatant was used in the assays. The dehalogenase activity assays (1 mL) were performed in capped 2 mL glass vials. Tris-HCl buffer (50 mM) was used for the assays at pH 6.0 and 7.0, and sodium acetate-acetic acid buffer (50 mM) was used for the assays at pH 4.75, 5.0, and 5.5. Ti(III) citrate-reduced methyl viologen (1 mM) was used as the artificial electron donor and *cis*-DCE (0.5 mM) was used as the electron acceptor; these were added to the buffers before the addition of cell extracts. The cell extracts were added into the vials to a final protein concentration of 10 μg/mL to start the reaction. The vials were immediately capped, incubated for 2 h, and sampled for GC analysis. Protein concentration was determined using a microplate Bradford (Sigma –Aldrich Bradford Reagent) protein assay with bovine serum albumin standards.

### *Sporomusa* growth inhibition assays

For this experiment, Na_2_S (0.2 mM), instead of FeS, was used as the reductant in the medium to avoid turbidity and facilitate cell density measurements; resazurin was excluded from the medium to avoid interference from the pink color that develops during cell density measurements after exposure to air. In an anaerobic chamber, the active *Sporomusa* sp. strain KB-1 pure culture (2 mL; OD_600nm_ ~ 1), was transferred into 2-mL sterile O-ring-capped plastic microcentrifuge tubes and pelleted down by centrifugation at 10,000 x g for 10 min at room temperature. The supernatants were removed, and the pellets were washed with sterile cobalamin-free defined medium (1 mL) twice. After, the pellets were resuspended in sterile defined media containing varying cobalamin concentrations (0, 5, 10, 25, 50 μg/L) to a final cell density (OD_600 nm_) of 0.04 (as determined by the no-substrate control included in the experiment); 5 mL of the resuspended cells at each cobalamin concentration were transferred into the sterile 10-mL rubber stopper-sealed glass tubes by sterile plastic syringes and needles. The tubes were then purged with 20% CO_2_/80%N_2_ to remove H_2_, followed by the addition of methanol to a final concentration of 20 mM. In some assays, H_2_ (20 % in head space), formate (10 mM), or lactate (10 mM) was added. The glass tubes were incubated at 30°C for one day, and 0.2 mL of each culture was sampled and loaded into a 96-well optical plate (Falcon). The cell density in each well was determined using a TECAN Infinite M200 plate reader at 600 nm.

## RESULTS AND DISCUSSION

### Reductive dechlorination of TCE to ethene in enrichment cultures grown without exogenous vitamins

Dechlorination of TCE to ethene was sustained over 800 days (Figure 1) in all the cultures, TCE/M_Vit(+), TCE/M_B12(−), and TCE/M_Vit(−), regardless of the type of vitamin amendment. In general, the cultures with or without exogenous vitamins performed similarly (Figure 1A and 1C). The average long-term dechlorination rates for these three cultures, taken as the slope of the linear regression of the data over 800 days as shown in Figure 1, were 0.048 ± 0.002, 0.046 ± 0.002, and 0.040 ± 0.001 mmol Cl^−^ released/L/d, respectively. These long-term data demonstrate that the entire vitamin requirements of the culture could be met by organisms present in the KB-1 culture.

**Figure 1.**
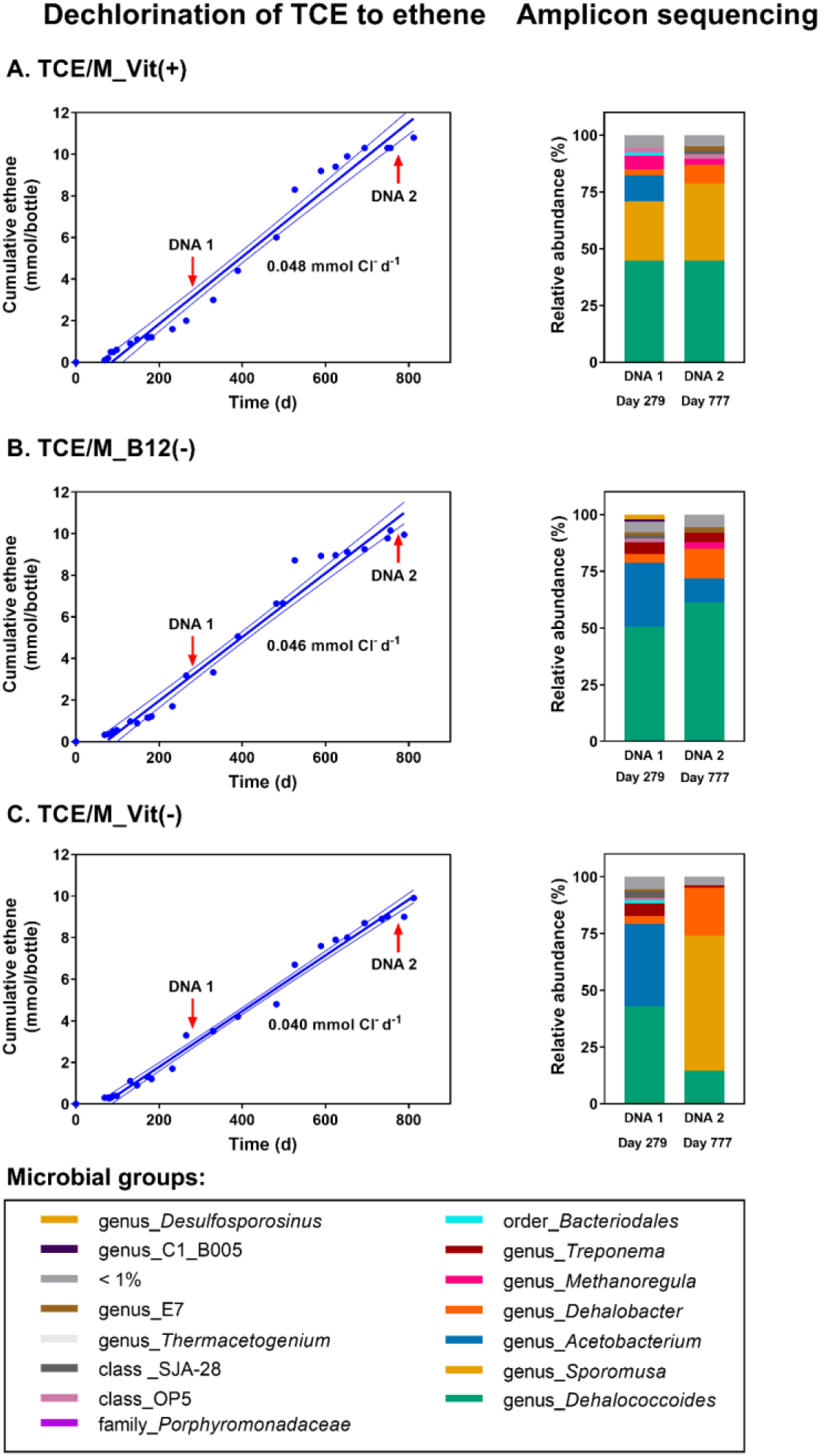
Long-term and sustained dechlorination of TCE to ethene (left) and corresponding microbial community composition shifts (right) for the scaled-up TCE- and methanol-fed enrichment cultures TCE/M_Vit(+) (A), TCE/M_Vit(−) (B), and TCE/M_B12(−) (C). TCE was fed every 4 to 6 weeks, at a nominal concentration of 1.4 to 1.5 mM. Dechlorination rates obtained from linear regressions are shown with 95% confidence intervals; the R^2^ values of the regressions are 0.98, 0.99, and 0.98, for (A), (B) and (C), respectively. On the right, two amplicon sequencing events are shown for DNA extracted on Day 279 and Day 777.

### Reductive dechlorination of VC to ethene in enrichment cultures grown at pH 6.0 and pH 5.5

The reductive dechlorination of VC to ethene in the cultures VC/M_pH7, VC/M_pH6, and VC/M_pH5.5 is shown in Figure 2A. The observed dechlorination rates for these cultures, taken as the slope of the linear regression of the data set shown for each culture (~ 400 days), are 7.9 ± 0.29, 8.8 ± 0.25, and 2.0 ± 0.08 μmol Cl^−^ released/L/d, respectively. The enrichment culture originally set at pH 6.0, described in the Master’s thesis of Li (2012)^30^, initially dechlorinated VC more slowly than the culture at pH 7.0, but eventually maintained a rate comparable to that of the enrichment culture maintained at pH 7.0 (Figure 2). In the culture maintained at pH 5.5 (see Figure S3 for pH measurements), the observed dechlorination and ethene production rates were about 4 times lower than those at pH 7.0 and 6.0. The maintenance of low pH batch cultures required careful monitoring as acid build up from dechlorination progressively dropped the pH; at low pH, the carbonate buffering system is not as robust as at neutral pH. Overall, we found that the enrichment culture KB-1 was able to dechlorinate VC to ethene at 5.5 ≤ pH ≤ 7.0 (Figure 2A), and that this activity was mediated by *Dehalococcoides* (Figure 2B), which is different from the observations reported by Yang et al.^24^ Perhaps, certain *Dehaloccocoides* populations are more resistant to low pH than others.

**Figure 2.**
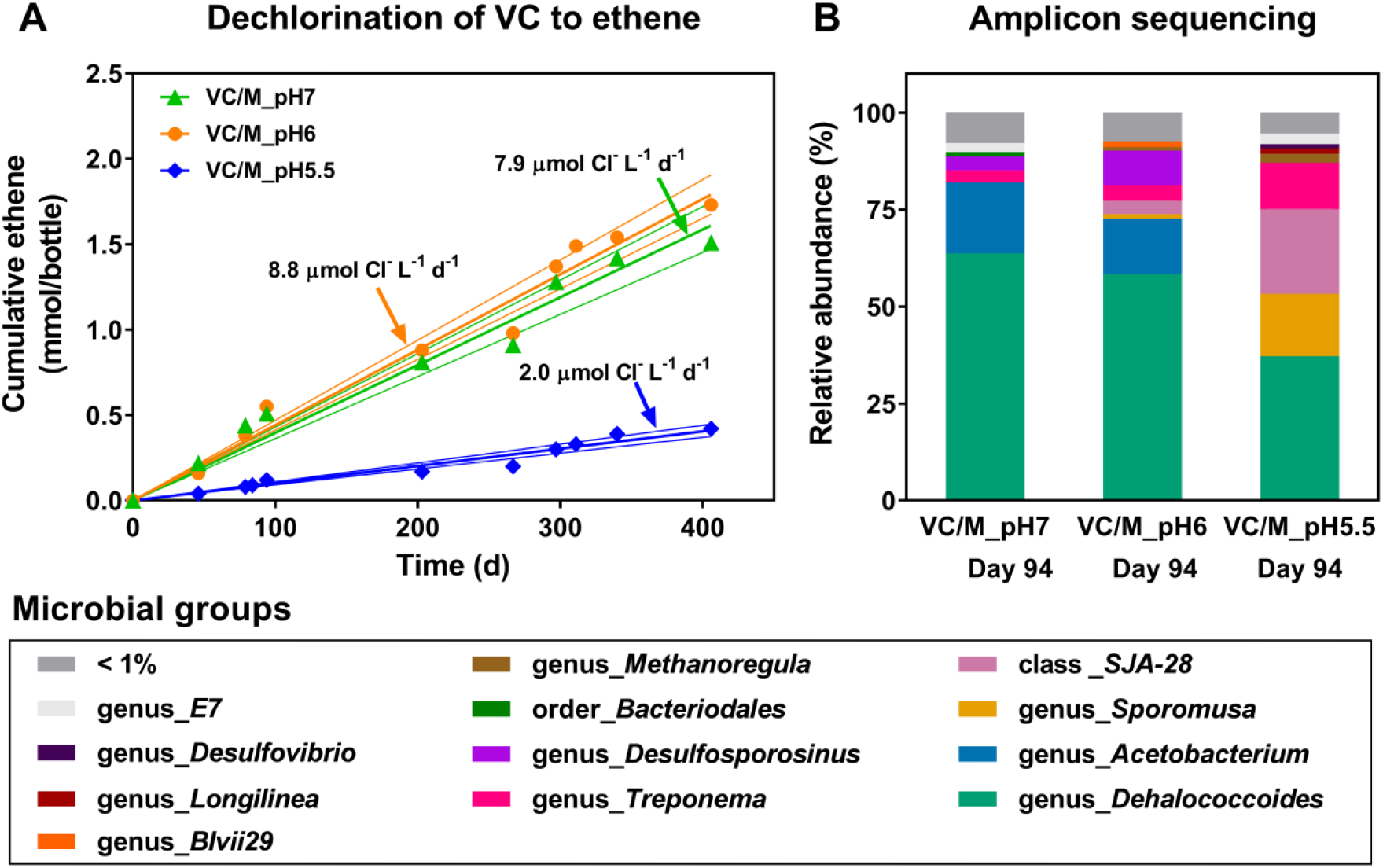
Long-term and sustained dechlorination of VC to ethene in enrichment cultures maintained at pH 7.0, 6.0, and 5.5 ± 0.2 (A), and their corresponding microbial community composition on Day 94 (B). Dechlorination rates obtained from linear regressions with 95% confidence intervals are shown; the R^2^ values of the linear regressions are 0.96, 0.97, and 0.96, for the pH 7.0, pH 6.0, and pH 5.5 enrichment cultures, respectively.

### Microbial community composition revealed by amplicon sequencing

Two snapshots of the microbial community composition in the enrichment cultures used in this study are shown in the right-hand panels in Figures 1 and 2. As expected, *Dehalococcoides* was the dominant genus in these cultures, accounting for about 50% of the total microbial population based on 16S rRNA gene copies. As previously reported^3, 17, 18^, the two major acetogens were *Acetobacterium* and *Sporomusa*. *Sporumusa* was more abundant in the enrichments amended with vitamins (TCE/M_Vit(+)) than in the non-amended ones. *Sporomusa* was also more abundant than *Acetobacterium* in the sample from VC/M_pH5.5. The relative abundance of *Sporomusa* increased in the TCE/M_Vit(−) enrichment cultures between Day 279 and Day 777. We also noted that the pH in this culture and in the TCE/M_Vit(+) had fallen to 4.7-4.8 (as a result of progressive HCl accumulation from dechlorination). The pH was promptly adjusted back to 7.0, but perhaps this pH drop had already affected the microbial community. Nevertheless, these observed differences in the relative abundance of *Acetobacterium* and *Sporomusa* led us to propose two hypotheses: 1) *Sporomusa* outcompetes *Acetobacterium* at pH below pH 6.0; and 2) *Sporomusa* outcompetes *Acetobacterium* when exogenous vitamins (particularly vitamin B_12_) are present. To test the pH hypothesis further, we carried out a point biserial (pb) correlation analysis between the microbial abundance data and pH above or below 6.0. We used the data shown in Figures 1 and 2, plus two samples from the parent KB-1 culture and a VC/methanol-fed KB-1 subculture (see Table S2). The relative abundance of *Acetobacterium* was negatively correlated (r_pb_ = −0.75) with pH<6.0 (anytime within three weeks prior to DNA extraction). In contrast, the relative abundance of *Sporomusa* was positively correlated (r_pb_ = 0.77) with pH>6.0. *Acetobacterium* also seemed to be more abundant in cultures without added vitamin B_12_ (Figures 1B and 1C). Additional growth experiments (described below) were initiated to further investigate these trends.

### Dechlorination of TCE at pH 7.0 and pH 5.5 in cultures grown without exogenous cobalamin (vitamin B_12_)

Using a 10% inoculum from culture TCE/M_B12(−), we examined the growth of *Dehalococcoides, Acetobacterium*, and *Sporomusa* (measured by qPCR) during the dechlorination of TCE to ethene in parallel experiments at pH 7.0 and 5.5. Gene copies per mL were converted to cells per mL using the number of 16S rRNA gene copies found in the genomes of *Acetobacterium* sp. KB-1 (5 copies, draft genome in NCBI under accession number CP030040) and *Sporomusa* sp. KB-1 (13 copies, draft genome in IMG under genome ID: 2512047088). *Dehalococcoides* only possess a single copy of the 16S rRNA gene per genome. As shown in Figures 3A and 3B, complete dechlorination of TCE to ethene was only observed at pH 7.0. At pH 5.5, TCE was dechlorinated to VC, but VC was not further dechlorinated to ethene. In these bottles, the dechlorination of TCE (~ 0.75 mM per bottle) to VC at pH 5.5 caused the pH to drop to 5.0. This drop in pH likely impacted the ability of the *Dehalococcoides* population to dechlorinate VC to ethene. Nonetheless, an increase in the cell numbers of *Dehalococcoides* per mL of culture (as compared to the measurements obtained at t = 0 d) was observed at both pH 7 and 5.5 (Figures 3C and 3D; Table S3); at pH 5.5, the growth of *Dehalococcoides* was proportional to observed dechlorination (two thirds of the electron acceptor equivalents were consumed at pH 5.5 compared to pH 7). In the transfer set up at pH 7.0, there was a 5-fold increase of *Acetobacterium*, but no increase of *Sporomusa*. In the transfers set up at pH 5.5, *Acetobacterium* and *Sporomusa* both showed a small 2-fold increase (Figures 3C and 3D; Table S3). Slow to no growth of *Acetobacterium* at pH 5.5 was expected as the pH growth optimum for other *Acetobacterium* spp. was previously shown to be between 7.0 and 8.0.^32–34^

**Figure 3.**
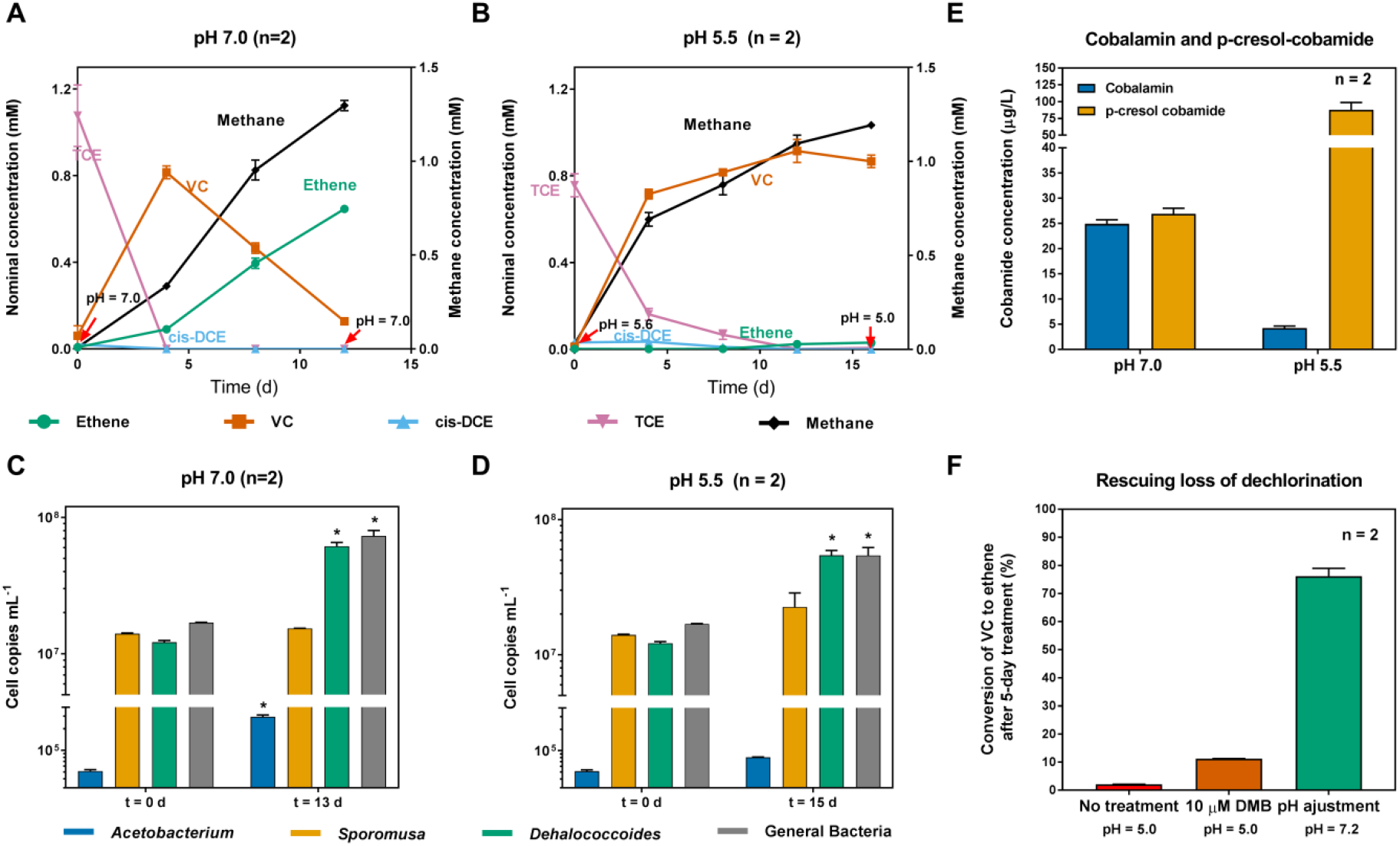
TCE dechlorination profiles in 10% dilution transfers of the TCE/M_B12(−) at pH 7.0 (A) and pH 5.5 (B), and associated cell and corrinoid abundances. Corresponding cell abundances per mL of culture as determined by qPCR are shown for pH 7.0 in panel C and pH 5.5 in panel D. Cobalamin and p-cresol-cobamide concentrations are shown in panel E. Attempts to reactivate pH 5.0 culture samples from Day 16 in Panel B with DMB addition or pH adjustment are shown in Panel F. In panels A and B, initial and final pH measurements are shown. Error bars correspond to the range of duplicate experimental bottles except for the qPCR samples at t = 0 d in which a single sample was measured and error bars correspond to the range for technical duplicates. The * symbol indicates growth greater than five-fold as compared to the measurements obtained at t = 0 d. In E, cobalamin and p-cresol-cobamide were measured at t = 16 d (final time point for the transfers set up at pH 5.5).

### Cobamide profile in cultures grown at pH 7.0 and 5.5

Since *Acetobacterium* spp. are reported to be cobalamin producers^13, 14^, we hypothesized that the amount of cobalamin in the cultures should be proportional to the relative abundance of the *Acetobacterium* population. Thus, we extracted the cobamides from the aforementioned KB-1 transfers grown at pH 7.0 and pH 5.5 (Figure 3E; MS spectra in Figure S4). Consistently, in the cultures grown at pH 5.5 in which VC accumulated, the measured concentration of p-cresol-cobamide was ~88 μg/L which is ~20 times higher than the measured concentration of cobalamin (~4 μg/L). In contrast, in the cultures grown at pH 7.0, in which VC was completely dechlorinated and the growth of *Acetobacterium* was measureable (5-fold increase), the average concentration of p-cresol-cobamide and cobalamin are comparable (~25 μg/L). The higher cell density of *Acetobacterium* and higher cobalamin concentration in the cultures grown at pH 7.0, as compared to the cultures grown at pH 5.5, supports our hypothesis that *Acetobacterium* is a DMB/cobalamin producer in the KB-1 enrichment cultures. To further confirm that *Acetobacterium* sp. strain KB-1 can supply DMB/cobalamin to *Dehalococcoides*, we inoculated medium without cobalamin with a 1% transfer from the culture TCE/M_Vit(−). In the previous experiment (10% dilution transfers), some DMB or cobalamin could have carried over with the inoculum. For this experiment, the inoculum was centrifuged and washed twice with anaerobic mineral medium devoid of vitamins to remove cobalamin and DMB carryover. After the duplicate KB-1 cultures had dechlorinated ~0.5 mM of TCE to ethene (1.34 mM Cl^−^ released), we harvested one of the cultures (dechlorination data shown in Figure S5) and measured cobalamin concentrations. The measured concentration (~750 ng/L, 0.55 nM) is almost 20-fold higher than the observed concentration before TCE dechlorination occurred (Table 1). Nevertheless, it is surprising that nanomolar levels of cobalamin could support the dechlorination of TCE to ethene. For this experiment, the yield of *Dehalococcoides*, 1.1 x 10^8^ cells per μmol Cl^−^ released, is in the same order of magnitude as previously published yields of *Dehalococcoides* pure cultures which range between 1.5 x 10^7^ to 2.9 x 10^8^ cells per μmol Cl^−^ released.^35^ Moreover, the cell density for *Acetobacterium* increased by 63-fold (Table 1).

**Table 1.** Cell growth, yield, and cobalamin production measured in a 1% dilution transfer of the TCE/M_Vit(−) enrichment culture dechlorinating TCE to ethene in the absence of any added vitamins.

| Measurement | Inoculum (t = 0 d) | Final (t = 60 d) | Yield (cells per $\mu\text{mol Cl}^-$ released) | Fold change |
| --- | --- | --- | --- | --- |
| *Dehalococcoides* (cells $\text{mL}^{-1}$) | $2.4 \pm 0.4 \times 10^7$ | $1.7 \pm 0.05 \times 10^8$ | $1.1 \pm 0.04 \times 10^8$ | **7.3** |
| *Acetobacterium* (cells $\text{mL}^{-1}$) | $5.7 \pm 0.2 \times 10^3$ | $3.6 \pm 0.04 \times 10^5$ | - | **63** |
| Total Cobalamin (ng $\text{L}^{-1}$) | $40 \pm 5$ | $750 \pm 140$ | - | **19** |

Dechlorination data is shown in Figure S5. Cell growth and cobalamin concentrations are reported as the mean +/- range of duplicate measurements.

### The relationship between *Acetobacterium* and *Dehaloccocoides* during TCE reductive dechlorination

As demonstrated earlier, the growth of *Acetobacterium* in KB-1 is correlated to an increase in cobalamin content which supports efficient dechlorination of VC to ethene (Table 1). Analyses of the KB-1 metagenome (NCBI accession PRJNA376155) are consistent with this observation, revealing that only *Acetobacterium* sp. strain KB-1 contains a complete anaerobic DMB biosynthesis operon (*bza*ABCDE; Figure S6). The operon arrangement and gene annotation are identical to that of *Eubacterium limosum* strain ATCC 10825 and *Acetobacterium Woodii*, two well-known cobalamin producers^14, 36, 37^ (Figure S6); the encoded proteins are most similar to the ones found in *Acetobacterium dehalogenans* (Table S4).

We were next interested in studying the growth of *Acetobacterium* in the absence of dechlorination to further understand the relationship between *Acetobacterium* and *Dehalococcoides*. We performed a 10% dilution transfer of the culture TCE/M_B12(−) into medium free of vitamins amended with methanol but not TCE, at both pH 7.0 and 5.5. Contrary to our expectation, we did not observe growth of *Acetobacterium* (Figure S7). Measurable growth (an order of magnitude) was observed only for *Sporomusa*, regardless of pH. *Dehalococcoides* was not expected to grow at either pH tested, yet we expected to observe measurable growth for *Acetobacterium* analogous to the growth that we observed at pH 7.0 with added TCE (Figure 3C). While it is clear that DMB/cobalamin produced by *Acetobacterium* is beneficial to *Dehalococcoides*, any potential mechanism that may facilitate the growth of *Acetobacterium* during active dechlorination by *Dehalococcoides* remains to be further explored.

When cobalamin is absent at field sites, VC dechlorination by *Dehalococcoides* can be expected to be highly dependent on the biosynthesis of DMB/cobalamin by cobalamin-producing organisms such as *Acetobacterium*. The role that native *Acetobacterium* populations have at chlorinated ethene-contaminated sites should be investigated. Environmental practitioners working on biostimulation via cobalamin addition may find it beneficial to first assess if native cobalamin-producing populations may support in situ VC dechlorination. *Acetobacterium* can be quantified, via qPCR of the 16S rRNA gene for example, in addition to *Dehalococcoides* 16S rRNA genes, *Dehalococcoides* RDase genes, and 16S rRNA genes for other dechlorinating organisms. *Acetobacterium* may indeed serve as an additional biomarker for successful field site reductive dechlorination of VC.

### Cobalamin in methanol-fed enrichment cultures may inhibit *Sporomusa*

According to our findings, when enrichment cultures are starved of TCE (Figure S7) or experience low pH (Figures 2B and 3D), *Sporomusa* growth is favored. *Sporomusa* can also efficiently grow at neutral pH as shown in Figure S7. An earlier study^38^ reported that the addition of 10 μM DMB (~ 1400 μg/L) to growth medium inhibited the growth of *Sporomusa* on methanol since the methanol methyltransferase requires either phenyl-cobamide or p-cresol-cobamide as a cofactor. ^38–40^ We hypothesized that cobalamin, the cobamide with DMB as the lower ligand, may also inhibit the growth of *Sporomusa* on methanol. When 50 μg/L of cobalamin was added to the medium, the growth of *Sporomusa* was completely inhibited (Figure 4). When 25 μg/L of cobalamin was added to the medium, the growth of *Sporomusa* was lower relative to the control. This is about the same concentration as measured in the pH 7.0 experiment shown in Figure 3 in which the growth of *Sporomusa* was not apparent (Figure 3C). The growth of *Sporomusa* was not inhibited by cobalamin when grown with H2, formate, or lactate instead of methanol (Figure 4), suggesting that the inhibition by cobalamin is specific to growth on methanol which requires the p-cresol-cobamide- or phenyl cobamide-specific methanol methyltransferase. It is possible that the use of methanol as the sole electron donor in the KB-1 enrichment cultures may play a role in the enrichment of the DMB/cobalamin-producing *Acetobacterium* population, whose production of cobalamin represses the growth of the p-cresol-cobamide-producing *Sporomusa* population.

**Figure 4.**
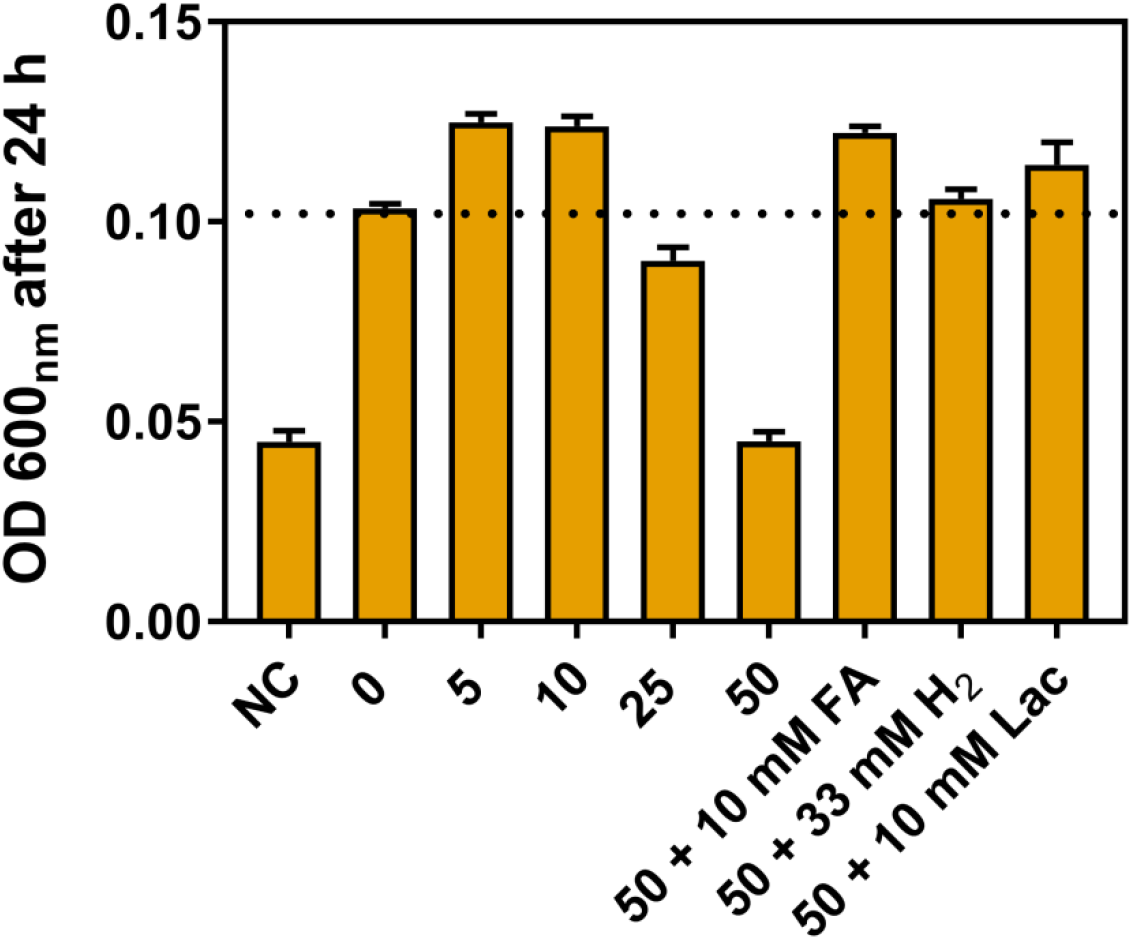
Inhibition of *Sporomusa* growth on methanol by cobalamin. Assays tested 0, 5, 10, 25, and 50 μg/L cobalamin plus 20 mM methanol; 50 μg/L cobalamin + 10 mM sodium formate (FA); 50 μg/L cobalamin + 33 mM (nominal) hydrogen gas; and 50 μg/L cobalamin + 10 mM sodium lactate (Lac). Growth was measured using OD_600nm_ after 24 hours. NC stands for the no-substrate control. Above the dotted line, no inhibition is observed.

### Stalled dechlorination of VC to ethene was restored after pH adjustment

Our data revealed that the population of cobalamin/DMB-producing *Acetobacterium* is much less abundant in the dilution transfer cultures grown at pH 5.5 than those grown at pH 7.0 (Figure 3), suggesting that *Dehalococcoides* may have faced cobalamin shortage to produce the functional form of VcrA for VC dechlorination. Given that *Dehalococcoides* can salvage cobalamin using p-cresol-cobamide and DMB,^12^ we wanted to test if DMB supplementation (10 μM; ~1400 μg/L) alone could rescue the loss of VC dechlorination in these transfers. As shown in Figure 3F, DMB supplementation did not restore the dechlorination of VC to ethene at pH 5.5. However, increasing pH to 7.0 led to 80% conversion of VC to ethene in 5 days. This suggests that incomplete VC dechlorination by *Dehalococcoides* was not due to DMB shortage but due to acidic pH.

### Effect of pH on the *Dehalococcoides-mediated* reductive dechlorination of VC in 10% dilution transfers amended with VC

Analogous to the 10% dilution transfer experiment set up at pH 7.0 and 5.5 with TCE as electron acceptor, a 10% dilution transfer experiment with the culture TCE/M_B12(−), maintained at circumneutral pH, was performed to specifically test the dechlorination of VC at different pH: 7.0, 6.3, 5.5, and 5.5 with 10 μM DMB. As shown in Figure 5A, complete VC dechlorination to ethene only occurred at pH 7.0 and pH 6.0. The dechlorination of VC to ethene was slower in the transfers set up at pH 5.5, but ethene was produced until the medium became acidic. *Dehalococcoides* growth was observed at each pH (Table 2). Even though the enrichment culture TCE/M_B12(−) is grown at circumneutral pH, *Dehalococcoides* was still able to grow and dechlorinate VC to ethene at a pH as low as 5.5. The addition of 10 μM of DMB at pH 5.5 resulted in enhanced *Dehalococcoides* growth (Table 2), 10 times (pH 5.5 + DMB) vs. 4 times (pH 5.5 without DMB addition), and a higher final pH, 4.8 (pH 5.5 + DMB) vs. 4.0 (pH 5.5 without DMB addition). Interestingly, DMB addition led to a noticeable pH buffering effect. In terms of yield, at pH 6.3 and 7.0, the yield of *Dehalococcoides* (1.1 ± 0.1 x 10^7^ cells per μmol Cl^−^ released) is in the same order of magnitude as previously published yields for *Dehalococcoides* pure cultures which range between 1.5 x 10^7^ to 2.9 x 10^8^ cells per μmol Cl^−^ released^41^, yet these published yields were measured in cultures generally grown with cyanocobalamin supplementation at 50 μg/L^2, 35^. The reported yields for the BAV1 and GT pure *Dehalococcoides* cultures grown with 1 μg/L of cyanocobalamin were 1.4 ± 0.8 x 10^7^ and 5.3 ± 1.3 x 10^7^ cells per μmol Cl^−^ release, respectively^12^, which are comparable to the yield obtained in this study.

**Figure 5.**
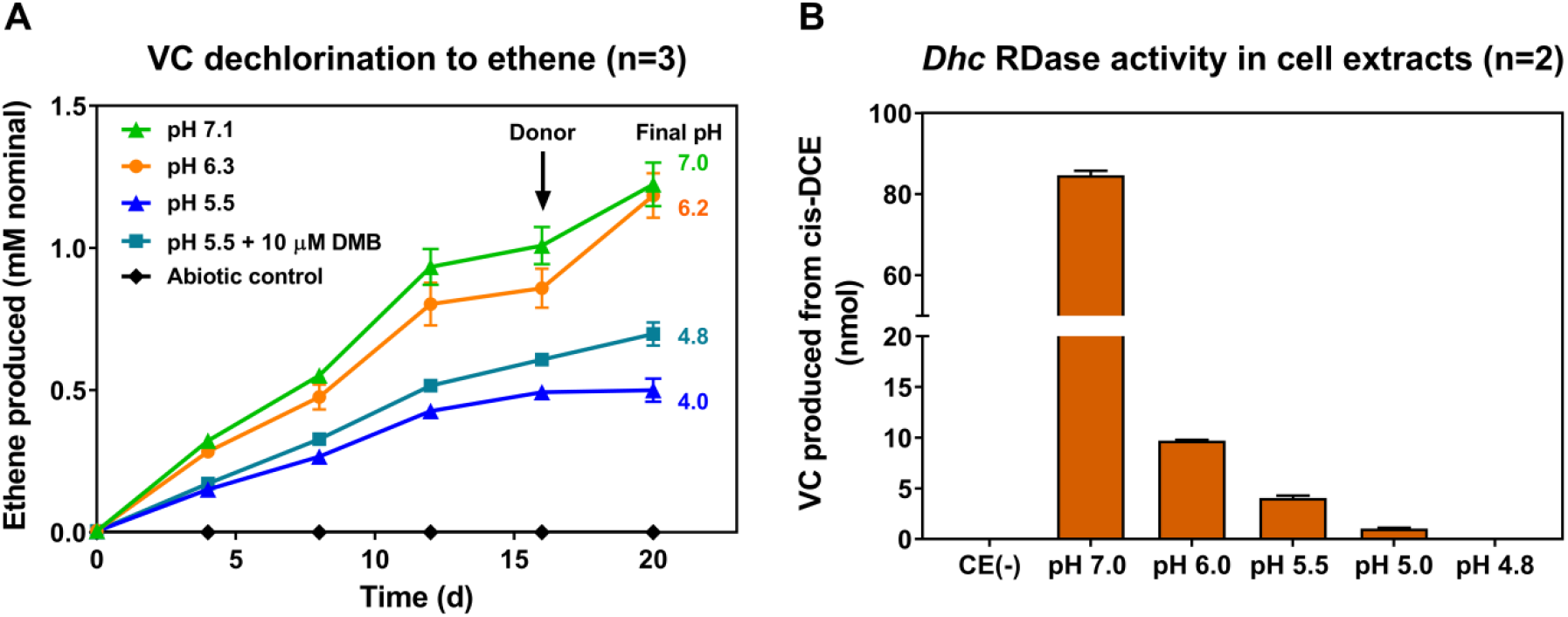
Effect of pH on the reductive dechlorination of VC by *Dehalococcoides* in KB-1 enrichment cultures originally grown at circumneutral pH. Dechlorination of VC to ethene in 10% dilution transfers of the TCE/M_B12(−) enrichment culture amended with VC and methanol (A) and *Dehalococcoides* (*Dhc*) RDase activity in cell extracts, amended with *cis*-DCE, of the TCE/M_Vit(−) enrichment culture, measured at varying pH (B). In A, the initial pH of the transfers is shown in the figure legend and the final measured pH is shown next to the last measured time point. In B, CE(−) stands for the control without cell extract. Error bars represent the standard deviation of triplicate experiments.

**Table 2.** Changes in *Dehalococcoides* cell numbers per mL in 10% dilution transfers of the TCE/M_B12(−) enrichment culture after the dechlorination of VC to ethene at varying pH.

| Starting pH | Final pH | *Dehalococcoides* cells $\text{mL}^{-1}$ | | $\text{Cl}^-$ released from VC (mmol) | Yield (cells per $\mu\text{mol Cl}^-$ released) | Fold change |
| --- | --- | --- | --- | --- | --- | --- |
|  |  | t = 0 d | t = 22 d |  |  |  |
|  |  | Mean | Mean |  |  |  |
| 5.5 | 4.0 | $7.1 \pm 0.3 \times 10^6$ | $2.8 \pm 2.8 \times 10^7$ | 0.23 | $4.5 \pm 6.1 \times 10^6$ | **4** |
| 5.5 + 10 $\mu\text{M}$ DMB | 4.8 | $7.1 \pm 0.3 \times 10^6$ | $6.9 \pm 0.6 \times 10^7$ | 0.32 | $9.7 \pm 0.9 \times 10^6$ | **10** |
| 6.3 | 6.2 | $5.0 \pm 0.5 \times 10^6$ | $1.2 \pm 0.1 \times 10^8$ | 0.54 | $1.1 \pm 0.1 \times 10^7$ | **25** |
| 7.1 | 7.0 | $2.6 \pm 0.6 \times 10^6$ | $1.3 \pm 0.2 \times 10^8$ | 0.55 | $1.1 \pm 0.1 \times 10^7$ | **50** |

The negative impact of low pH on reductive dechlorination has been well documented ^20–24, 42^. Although members of the genus *Sulfurospirillum* have been shown to reductively dechlorinate PCE and TCE to *cis*-DCE at low pH (pH 5.5),^24^ this is the first time that growth of *Dehalococcoides* populations has been demonstrated at pH 5.5 after a rapid decrease from pH 7.0 to 5.5 (Table 2), and also after progressive decreases and long-term cultivation at pH 5.5 (Figure 2). Thus, *Dehalococcoides* remains a key player in the complete chlorinated ethene detoxification at field sites even when pH is below 6.0.

### Dechlorination in cell extracts at different pH

To gain additional insight on the effect of pH on the *Dehalococcoides*-mediated reductive dechlorination, cell extracts, harvested from the TCE/M_Vit(−) enrichment culture while actively dechlorinating, were assayed at pH 4.75, 5.0, 5.5, 6.0, and 7.0. In this culture, the ratio of 16S rRNA gene copies to vinyl chloride reductase gene (*vcrA*) copies is about 1:1 with ~1 x 10^9^ copies per mL of culture (qPCR data shown in Figure S8). Titanium citrate-reduced methyl viologen and *cis*-DCE were used as the artificial electron donor and electron acceptor, respectively; *cis*-DCE was chosen as a suitable acceptor because *cis*-DCE can be more easily and reproducibly amended as opposed to VC which is a gas at room temperature. The dechlorination activity of the cell extracts declined sharply from pH 7.0 to 6.0 (Figure 5B). At pH < 5.0, the dechlorination activity was completely lost. These results are consistent with our previous findings in the 10% dilution transfer set up at pH 5.5: the pH dropped from 5.5 to 5.0 and VC dechlorination did not occur (Figure 3B). In accordance to the cell-extract assays, VcrA enzyme activity is not expected at such low pH. From earlier studies on the catalytic mechanism of RDases ^43, 44^, it was possible to infer that RDases would be affected at low pH, yet the functional pH range of *Dehalococcoides* RDases had not been previously studied.

### Implications for environmental science and remediation

Vitamins are key molecules for various microbially-mediated processes and only a concentration of micrograms per litre is generally required. However, microbial vitamin biosynthesis has a high metabolic cost, *e.g*., more than 30 genes are involved in the anaerobic biosynthesis pathway of cobalamin.^45^ Thus, mixed microbial bioaugmentation cultures that do not require vitamin amendments are highly desirable at field sites to avoid the need for adding costly vitamins. Here, we have demonstrated that complete dechlorination of TCE to ethene occurs steadily in the KB-1 enrichment cultures grown without any exogenous vitamins (Figure 1). Vitamin autotrophy in mixed microbial communities is possible if the necessary vitamin-producing populations are present and maintained during cultivation. In KB-1, these vitamin-producing populations have been maintained owing to batch cultivation and infrequent medium changes. Dechlorinating cultures that grow in the absence of cobalamin have been reported before, yet incomplete dechlorination and VC accumulation occurred. ^46, 47^ To our knowledge, this is the first report of complete and sustained dechlorination to ethene in dechlorinating mixed cultures grown without any exogenous vitamins.

In a mixed microbial dechlorinating culture, and likely at field sites in which reductive dechlorination occurs, vitamin availability and pH impact the rate and extent of dechlorination. Our results indicate that slow dechlorination rates at low pH (<5.5) are mainly due to the loss of function of *Dehalococcoides* RDases. The activity of *Dehalococcoides* RDases in cell-extracts is completely lost below pH 5.0 (Figure 5B). Clearly, a pH above 5.0 is essential for the successful *Dehalococcoides-mediated* bioremediation of vinyl chloride (VC) at contaminated field sites regardless of the availability of DMB/cobalamin.

## ACKNOWLEDGEMENTS

We would like to thank Dr. Kirill Krivushin for his earlier work regarding the dominant acetogens, *Sporomusa* and *Acetobacterium*, in the KB-1 culture. We also thank Professor Frank E. Löffler and Dr. Jun Yan at the University of Tennessee, Knoxville for maintaining and supplying the pure culture of *Sporomusa* sp. strain KB-1 as well as providing us with the p-cresol-cobamide standard. We are grateful to Nadia Morson (U of Toronto) for performing the qPCR assays to determine the copy numbers of the vinyl chloride reductase (*vcrA*) gene in the TCE/M_Vit(−) culture, and to Robert Flick at the BioZone Mass Spectrometry facility (U of Toronto) for assisting with the UPLC-ESI-HRMS analyses.

## SUPPORTING INFORMATION

Supplementary tables, figures and detailed analytical procedures (PDF)

