## Supplementary Information for "Sustained dechlorination of vinyl chloride to ethene in *Dehalococcoides*-enriched cultures grown without addition of exogenous vitamins and at low pH"

### Supplementary Materials and Methods

**Culture pH monitoring and adjustments.** During culture maintenance and experiment set up, pH was monitored using an Oakton pHTestr device calibrated daily with three standard solutions at pH: 4.0, 7.0, and 10.0. Adjustments in pH were performed using a saturated sodium bicarbonate solution (~1.3 M) or 5N HCl, unless otherwise stated. For each target pH, a deviation of ± 0.2 units was considered acceptable.

**Analytical methods.** Chloroethenes, ethene, and methane were quantified using gas chromatography with flame ionization detection (GC-FID). Calibration standards were prepared gravimetrically in water using stocks prepared in methanol. Liquid culture samples (1 mL) were added to 5 mL of acidified (using 6N HCl) deionized water (pH < 2). The samples were equilibrated in an Agilent G1888 autosampler at 70°C for 40 min. After equilibration, headspace samples (3 mL) were automatically injected into an Agilent 7890A GC equipped with an Agilent GS-Q plot column (30 m length, 0.53 mm diameter) via a packed inlet. Helium was used as the carrier gas at a flow rate of 11 mL $min^{-1}$. The temperature of the injector and the detector were set at 200°C and 250°C, respectively. The oven was held at 35 °C for 1.5 min, ramped to 100°C at a rate of 15°C per min, ramped to 185°C at a rate of 5°C per min, held for 10 min, ramped to 200°C at a rate of 20°C per min, and held at 200°C for 10 min.

**DNA sampling and quantitative PCR (qPCR).** DNA was extracted from 2 mL liquid culture samples. Cells were harvested by centrifugation at 10,000 x g for 10 min at 4°C. The cell pellet was resuspended in 100 µl of supernatant and the DNA was extracted using the MO BIO PowerSoil® DNA isolation kit following the manufacturer's recommendations. Real-time polymerase chain reaction (qPCR) assays were performed to track the gene copy numbers of total bacteria, *Dehalococcoides*, *Acetobacterium*, and *Sporomusa* using specific 16S rRNA gene primers (see supplementary Table S1 for primer sequences). The qPCR prep was conducted in a UV-treated PCR cabinet (ESCO Technologies, Hatboro, PA) with the fan off. Each reaction mixture (20 µl) contained 10 µl of 2×SsoFast™ EvaGreen® (Bio-Rad, Hercules, CA), forward and reverse primers (0.5 µM each), and 2 µl of template DNA. The amplification program included an initial denaturation step at 98°C for 2 min, followed by 39 cycles of 5 s at 98°C and 10 s at the corresponding annealing temperature of each primer set. Quantification was performed using 10-fold serial dilutions of plasmid DNA as standards. The analyses were conducted using a BIO-RAD CFX96 Touch™ Real-Time PCR Detection System and the CFX Manager software. The number of gene copies per mL of culture was calculated assuming a 100% DNA extraction efficiency and taking into account the DNA dilution and culture volumes used for the extraction. The large variability of 16S rRNA gene copy numbers in bacterial genomes has been well documented.[30] In our study, for the purpose of calculating accurate cell numbers, the gene copies per mL of culture were divided by the number of 16S rRNA gene copies: 5 copies for *Acetobacterium* sp. KB-1 (draft genome in NCBI under accession number CP030040) and 13 copies for *Sporomusa* sp. KB-1 (draft genome in IMG under genome ID: 2512047088). For *Dehalococcoides* spp., only single copies of the 16S rRNA gene have been reported.

**16S rRNA gene amplicon sequencing.** DNA was extracted as described above and amplified via PCR using the universal primer set 926f (5′-AAACTYAAAKGAATTGACGG-3′) and 1392r (5′-ACGGGCGGTGTGTRC-3′), which targets the V6−V8 variable region of the 16S rRNA gene from bacteria and archaea. The PCR products were checked on a 2% agarose gel and replicate reactions were combined and purified using the GeneJET™ PCR Purification Kit (Fermentas, Burlington, ON), according to the instructions of the manufacturer. A NanoDrop ND-1000 Spectrophotometer was used to obtain approximate DNA concentrations. The cleaned PCR products were checked on a 2% agarose gel, and their concentrations were also compared to known concentrations of serial dilutions of DNA ladder. Samples were sent to the Genome Quebec Innovation Centre (McGill University) for amplicon sequencing using the Roche GS FLX Titanium technology (Roche Diagnostics Corporation, Indianapolis, IN). After sequencing, the raw data was processed using the QIIME package[31] v1.8.0 with default parameters unless otherwise noted below. The sequence data was demultiplexed and filtered using the following cut-offs: 250 bp minimum length and 8 bp maximum homopolymer length. Chimera checking and OTU picking were performed using usearch61[32, 33] v6.1.544. The seed sequence of each cluster was used as the representative OTU sequence. Taxonomy was assigned using the RDP classifier[34] v2.2 and the Greengenes reference database[35] version 13_8 clustered at 97% identity.

**Cobamide extraction.** Cobamides were extracted and derivatized following previous established protocols with some modifications. [37, 38] For the extractions performed with the 10% dilution transfers set up at pH 7.0 and 5.5, amended with TCE (see Figure S1), cells were harvested and pelleted down via centrifugation at 9,000 x g for 20 min at room temperature. For the extractions performed with the 1% dilution transfer, cells were harvested by filtration into 0.22 µm sterivex filters. The cell-containing filter papers were cut and placed in 50 mL falcon tubes for cobamide extraction. The pellets or cell-containing filter papers were resuspended in a solution of 80% methanol containing 10 mM KCN, and the pH was adjusted with 3% acetic acid (v/v) to a pH ranging between 5.0 to 5.5. Then, they were transferred into 2 mL plastic O-ring-capped microcentrifuge tubes and placed in a 80°C water bath for 60 min; the tubes were vortexed briefly every 15 min. After cooling down to room temperature, the tubes were centrifuged at 13,000 x g for 10 min to remove cell debris. The supernatants were transferred to the 2 mL plastic microcentrifuge tubes, incubated in a rotary evaporator until dried, and resuspended by 0.1 mL Milli-Q water for LC-MS analysis.

**Cobamide measurements.** Cobamides were quantified using liquid chromatography coupled mass spectrometry (LC-MS) with an Accela HPLC system and a Q-Exactive mass spectrometer equipped with HESI II sources (Thermo Scientific). System control and data handling were performed using the software Thermo XCalibur 2.2. Separation by liquid chromatography was conducted on a Hypersil Gold C-18 column (50mm x 2.1 mm, 1.9 µm particle size, Thermo Scientific) equipped with a guard column. LC was performed with 10 µL injections at a flow rate of 0.2 ml/min with a gradient of water containing 2.5 mM ammonium acetate (pH 6.0) (A) and methanol (B), and a column temperature of 30˚C. The mobile phase composition began at 100% A for 0.81 min, followed by a linear gradient to 15% B at 3.32 min, 50% B at 4.86 min, 90% B at 5.56 min, and 0% B at 5.59 min, followed by equilibration for 5 min at starting conditions prior to the next run. Cobamide data collection was done in positive ionization mode with a *m/z* scan range of 1000-1500; resolution 100,000 at 1 Hz, automatic gain control (AGC) target of 5e5; and a maximum injection time of 200 ms. The concentration of cobamide in the extracts was determined using external calibration curves generated with cobamide standards (cyanocobalamin and cyano-p-cresol-cobamide).

Table S1. Quantitative PCR (qPCR) primer sequences used in this study.

| Target genus | Primer short name | Primer Sequence 5' - 3' | Reference |
|---|---|---|---|
| *Dehalococcoides* | Dhc 1f | GAT GAA CGC TAG CGG CG | Grostern and Edwards [1,] Hendrickson, et al. [2] |
| | Dhc 264r | CCT CTC AGA CCA GCT ACC GAT CGA A | |
| *Acetobacterium* | Aceto 527f | GGC TCA ACC GGT GAC ATG CA | Duhamel and Edwards [3] |
| | Aceto 784r | ACT GAG TCT CCC CAA CAC CT | |
| *Sporomusa* | Sporo 168f | TAG AGA TGG GTC TGC GTC TG | Duhamel and Edwards [3] |
| | Sporo 367r | TCG TCC CAA ACG ACA GAG CT | |
| General Bacteria | Bac 1055f | ATG GCT GTC GTC AGC T | Ferris, et al. [4], Amann, et al. [5] |
| | Bac 1392 | ACG GGC GGT GTG TAC | |

Table S2. List of samples from methanol-fed KB-1-derived enrichment cultures used for the point biserial correlation analysis of 16S rRNA gene amplicon sequencing data with pH.

| Culture name/ sample name | Electron acceptor | Growth medium | | | pH prior to DNA extraction | |
|---|---|---|---|---|---|---|
| | | Vitamins | Vitamins except B12 | No Vitamins | <6.0 | >6.0 |
| TCE/M_Parent | TCE | x | | | | x |
| VC/M_T3VC | VC | x | | | | x |
| TCE/M_Vit(+) Day 0 | TCE | x | | | | x |
| TCE/M_B12(-) Day 0 | TCE | | x | | | x |
| TCE/M_Vit(-) Day 0 | TCE | | | x | | x |
| TCE/M_Vit(+) Day 450 | TCE | x | | | x | x |
| TCE/M_B12(-) Day 450 | TCE | | x | | | |
| TCE/MVit(-) Day 450 | TCE | | | x | x | |
| VC/M_pH7 | VC | | x | | | x |
| VC/M_pH6 | VC | | x | | | x |
| VC/M_pH5.5 | VC | | x | | x | |

Table S3. Calibration information for the qPCR assays performed in this study.

| qPCR Target | Data included in | Slope | y-intercept | $R^2$ | Efficiency | Limit of detection[1] (copies $\mu L^{-1}$) | Limit of detection[1] (copies $mL^{-1}$ culture) |
|---|---|---|---|---|---|---|---|
| General Bacteria | Figure 3, Table 2 | -3.78 | 37.50 | 0.986 | 84% | 5.3E+02 | 1.3E+04 |
| *Dehalococcoides* | Figure 3, Table 2 | -3.54 | 36.97 | 0.997 | 92% | 5.3E+01 | 1.3E+03 |
| *Acetobacterium* | Figure 3, Table 2 | -3.57 | 37.47 | 0.997 | 91% | 4.1E+01 | 1.0E+03 |
| *Sporomusa* | Figure 3, Table 2 | -3.55 | 37.01 | 0.999 | 91% | 3.5E+01 | 8.7E+03 |
| General Bacteria | Figure S4 | -3.79 | 36.59 | 0.998 | 84% | 4.5E+02 | 1.1E+04 |
| *Dehalococcoides* | Figure S4 | -3.69 | 36.43 | 0.995 | 87% | 4.5E+01 | 1.1E+03 |
| *Acetobacterium* | Figure S4 | -3.36 | 33.72 | 0.999 | 99% | 7.4E+01 | 1.9E+03 |
| *Sporomusa* | Figure S4 | -3.58 | 35.05 | 0.999 | 90% | 3.5E+01 | 1.0E+03 |
| *Dehalococcoides* | Table 1 | -3.62 | 36.80 | 0.998 | 89% | 4.4E+01 | 1.1E+03 |
| *Acetobacterium* | Table 1 | -3.67 | 37.08 | 0.999 | 87% | 4.1E+01 | 1.0E+02 |

[1]The limit of detection (L.O.D.) was defined as the measured value of the lowest standard that is above any amplification observed in the blanks. For example, when no amplification was seen in the blank, the L.O.D. was in the order of $10^1$ gene copies $\mu l^{-1}$ of DNA. This translates to $10^3$ genes copies $ml^{-1}$ as shown below.

$$L.O.D. = 5.3 * 10^1 \frac{copies}{\mu l} * \frac{50\ \mu l\ of\ elution\ solution}{2\ ml\ of\ culture} = 1.3 * 10^3 \frac{copies}{ml\ of\ culture}$$

Table S4. Comparison between the proteins encoded by the bza operon of *Acetobacterium* sp. strain KB-1 and *Acetobacterium dehalogenans.*

| Putative gene | Closest protein match (as shown in the NCBI record) | Organism[1] | Pairwise amino acid identity (%) | NCBI accession number for closest protein match |
|---|---|---|---|---|
| **bzaA** | phosphomethylpyrimidine synthase | *Acetobacterium dehalogenans* | **95** | WP_026394015.1 |
| **bzaB** | B12 lower ligand biosynthesis ThiC-like protein BzaB | *Acetobacterium dehalogenans* | **94** | WP_026394014.1 |
| **cobT** | nicotinate-nucleotide--dimethylbenzimidazole phosphoribosyltransferase | *Acetobacterium dehalogenans* | **96** | WP_026394013.1 |
| **bzaC** | class I SAM-dependent methyltransferase | *Acetobacterium dehalogenans* | **87** | WP_084504898.1 |
| **bzaD** | B12 lower ligand biosynthesis radical SAM protein BzaD | *Acetobacterium dehalogenans* | **93** | WP_026394012.1 |
| **bzaE** | B12-binding domain-containing radical SAM protein | *Acetobacterium dehalogenans* | **91** | WP_026394011.1 |

[1]At the time of the analysis, using the blastp suite from NCBI, the proteins encoded by each of the genes in the bza operon of *Acetobacterium* sp. strain KB-1 (query sequences) had the highest pairwise amino acid identity to the ones found in *Acetobacterium dehalogenans*.

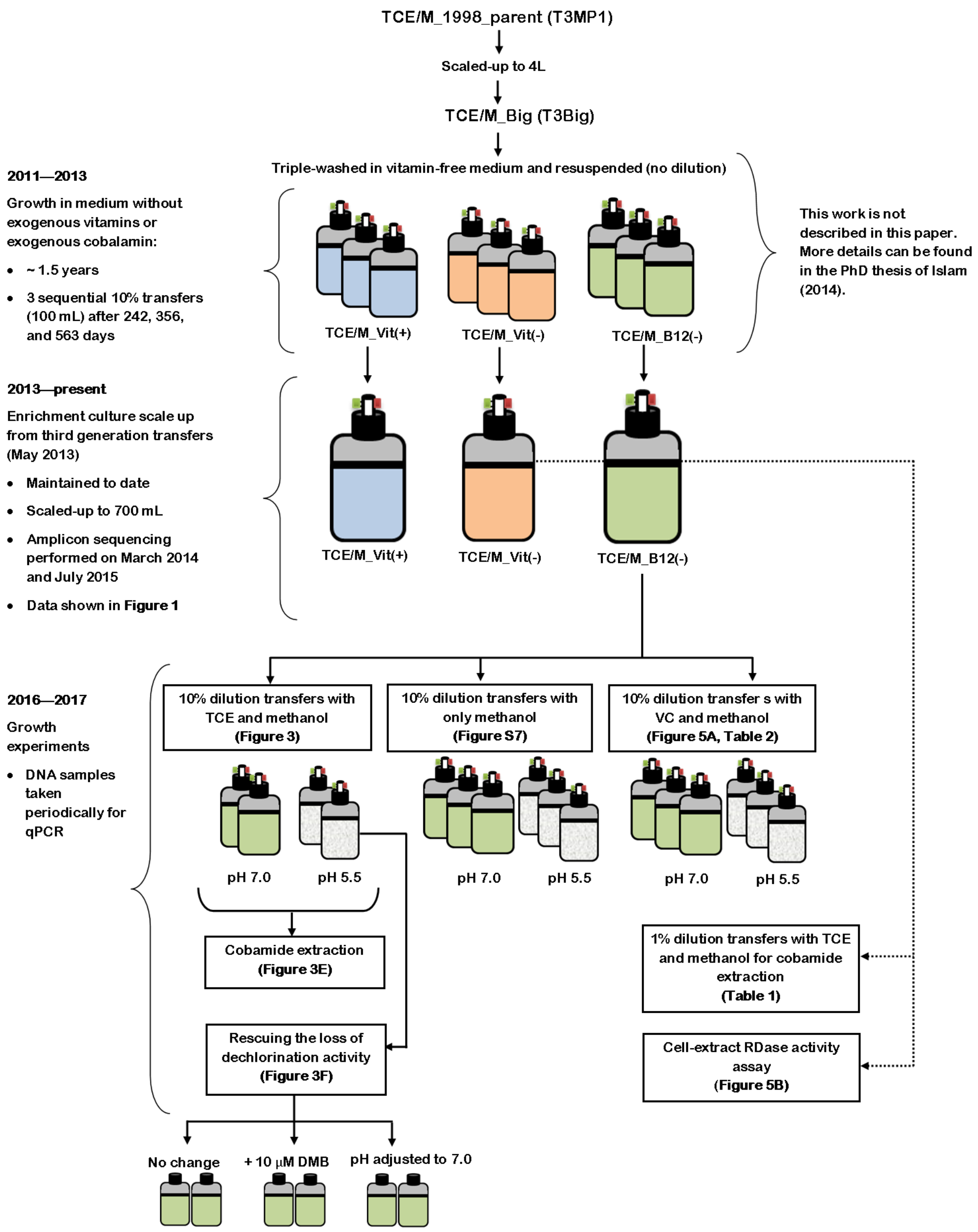

Figure S1. History and experiments performed with the enrichment cultures TCE/M_Vit(+), TCE/M_Vit(-), and TCE/M_B12(-).

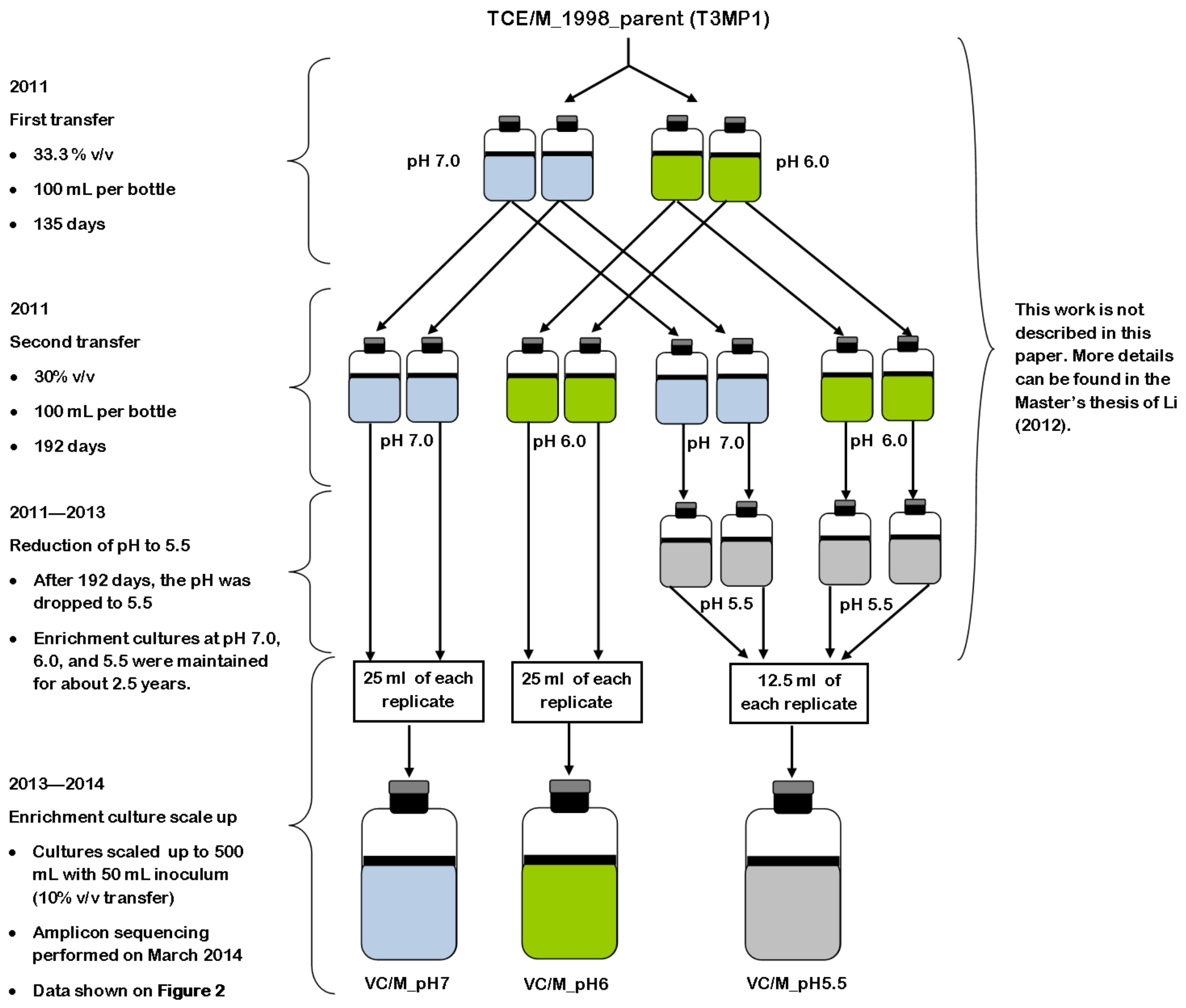

Figure S2. History of the VC and methanol-fed KB-1 enrichment cultures VC/M_pH7, VC/M_pH6, and VC/M_pH5.5 (see main Figure 2). All transfers depicted in the Figure were amended with VC and methanol.

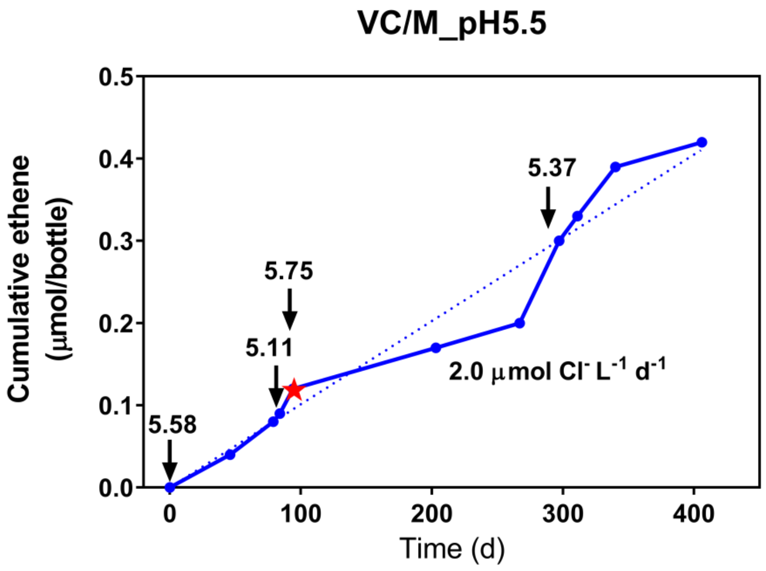

Figure S3. Cumulative ethene production in the VC/M_pH5.5 enrichment culture. Arrows indicate pH measurements. The red star indicates the time point of DNA extraction for 16S rRNA gene amplicon sequencing. This Figure complements the data shown in Figure 2. The data was collected between December of 2013 and January of 2015.

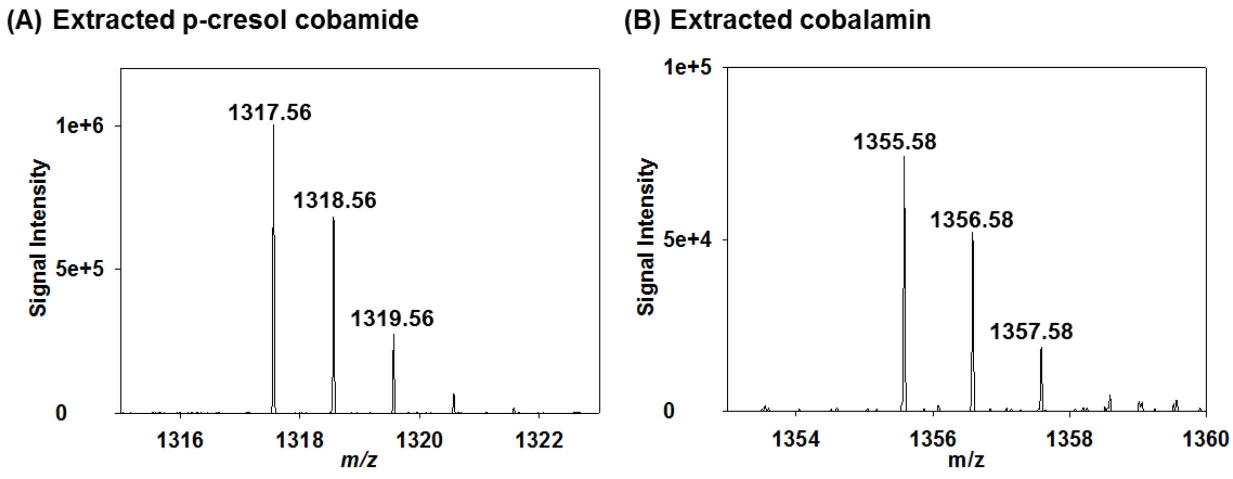

Figure S4. Cobalamin and p-cresol cobamide MS spectra (main data shown in Figure 3C).

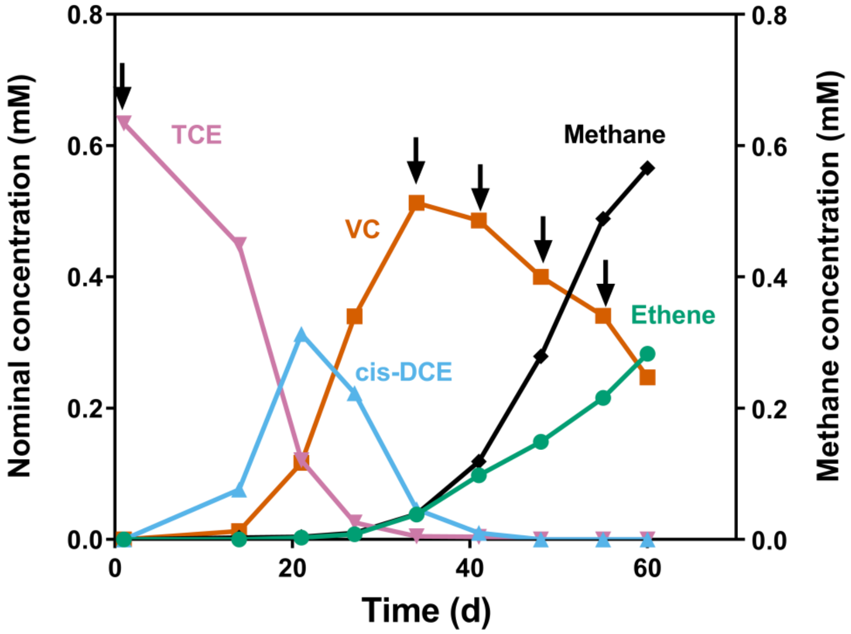

Figure S5. Dechlorination of TCE to ethene in a 1% dilution transfer, fed TCE and methanol, of the TCE/M_Vit(-) enrichment culture. Black arrows indicate methanol amendments. At t = 60 d, the culture was harvested for cobalamin extraction (see Table 1).

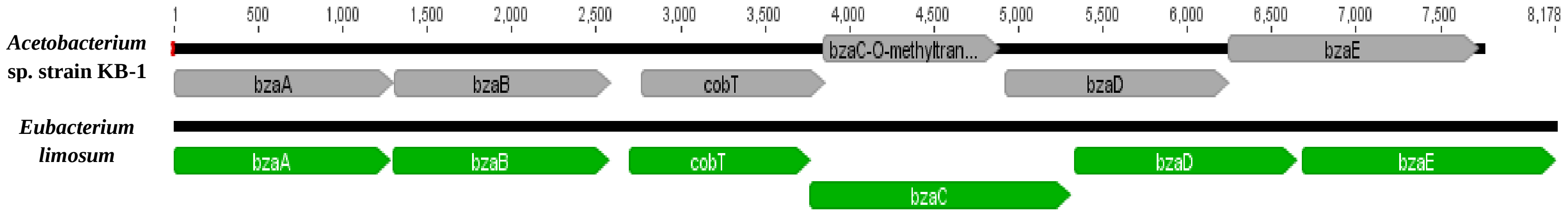

| Gene | bzaA | bzaB | cobT | bzaC | bzaD | bzaE |
|---|---|---|---|---|---|---|
| **Pairwise amino acid identity** | **76%** | **75%** | **71%** | **54%** | **69%** | **64%** |
| **SEED annotation in *Acetobacterium* sp. strain KB-1** | Hydroxy-methylpyrimidine phosphate synthase ThiC (EC 4.1.99.17) CDS | Hydroxy-methylpyrimidine phosphate synthase ThiC (EC 4.1.99.17) CDS | Nicotinate-nucleotide--dimethylbenzimidazole phosphoribosyltransferase (EC 2.4.2.21) CDS | O-methyltransferase, family 2 CDS | Radical SAM domain protein CDS | Radical SAM domain protein CDS |
| **NCBI annotation in *Eubacterium limosum*** | ThiC | ThiC | cobT | SAM methyltransferase superfamily protein | B12-binding-Fe-S/SAM superfamily protein | B12-binding-Fe-S/SAM superfamily protein |

Figure S6. DMB biosynthesis operon of *Acetobacterium* sp. strain KB-1 (grey) and characterized DMB biosynthesis operon (GI: 914703854) of *Eubacterium limosum* (green).

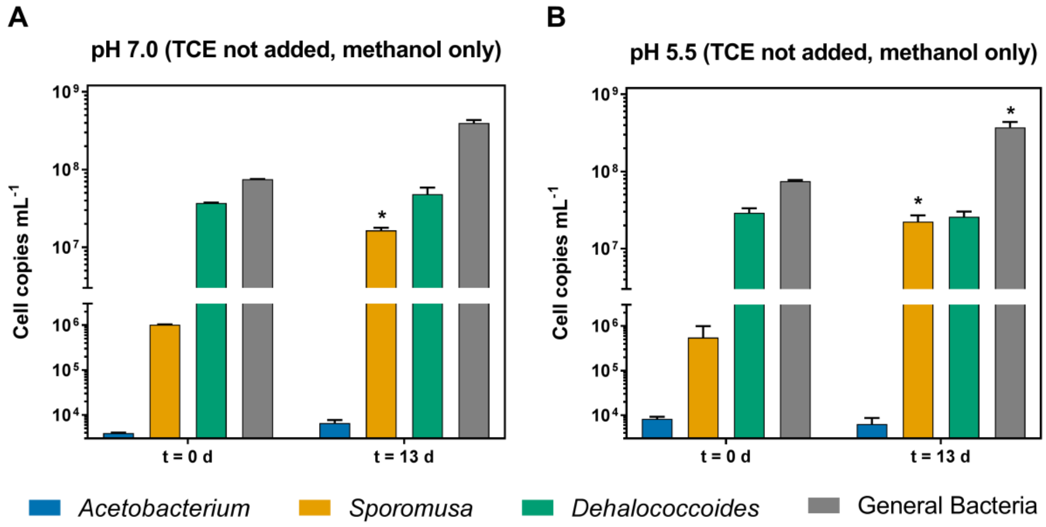

Figure S7. Cell copies per mL of culture as determined by qPCR of *Acetobacterium*, *Sporomusa*, *Dehalococcoides*, and General Bacteria, at pH 7.0 (A) and pH 5.5 (B) in 10% dilution transfers from the culture TCE/M_B12(-) when only methanol, and not TCE, was added to the growth medium. Error bars represent the range of duplicate qPCR assays (at t = 0 d) and standard deviation of triplicate experimental bottles (at t = 13 d). The * symbol indicates growth greater than five-fold as compared to the measurements obtained at t = 0 d

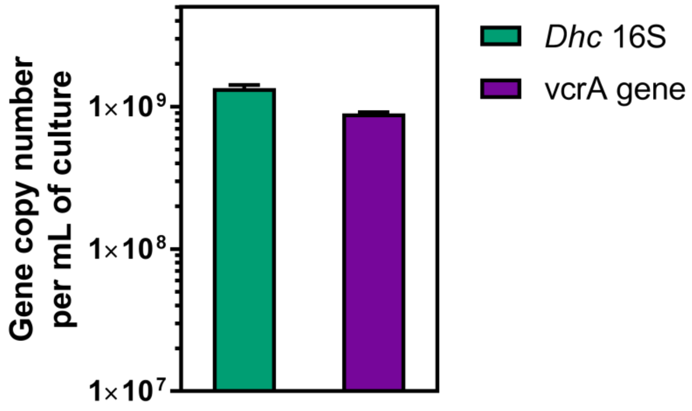

Figure S8. Gene copy number per mL of culture as determined by qPCR of the *Dehalococcoides* (*Dhc*) 16S rRNA and vcrA genes of the TCE/M_Vit(-) culture. Error bars represent the standard deviation of triplicate qPCR reactions (*Dhc* 16S) and the error of duplicate qPCR reactions
